# Evolution of andrenine bees reveals a long and complex history of faunal interchanges between the Americas during the Mesozoic and Cenozoic

**DOI:** 10.1101/2021.08.05.455338

**Authors:** Kelli S. Ramos, Aline C. Martins, Gabriel A. R. Melo

## Abstract

Bees are presumed to have arisen in the early to mid-Cretaceous coincident with the fragmentation of the southern continents and concurrently with the early diversification of the flowering plants. Among the main groups of bees, Andreninae sensu lato comprise about 3000 species widely distributed with greatest and disjunct diversity in arid areas of North America, South America, and the Palearctic region. Here, we present the first comprehensive dated phylogeny and historical biogeographic analysis for andrenine bees, including representatives of all currently recognized tribes. Our analyses rely on a dataset of 106 taxa and 7952 aligned nucleotide positions from one mitochondrial and six nuclear loci. Andreninae is strongly supported as a monophyletic group and the recovered phylogeny corroborates the commonly recognized clades for the group. Thus, we propose a revised tribal classification that is congruent with our phylogenetic results. The time-calibrated phylogeny and ancestral range reconstructions of Andreninae reveal a fascinating evolutionary history with Gondwana patterns that are unlike those observed in other subfamilies of bees. Andreninae arose in South America during the Late Cretaceous around 90 Million years ago (Ma) and the origin of tribes occurred through a relatively long time-window from this age to the Miocene. The early evolution of the main lineages took place in South America until the beginning of Paleocene with North American fauna origin from it and Palearctic from North America as results of multiple lineage interchanges between these areas by long-distance dispersal or hopping through landmass chains. Overall, our analyses provide strong evidence of amphitropical distributional pattern currently observed in Andreninae in the American continent as result at least three periods of possible land connections between the two American landmasses, much prior to the Panama Isthmus closure. The andrenine lineages reached the Palearctic region through four dispersal events from North America during the Eocene, late Oligocene and early Miocene, most probably via the Thulean Bridge. The few lineages with Afrotropical distribution likely originated from a Palearctic ancestral in the Miocene around 10 Ma when these regions were contiguous, and the Sahara Desert was mostly vegetated making feasible the passage by several organisms. Incursions of andrenine bees to North America and then onto the Old World are chronological congruent with distinct periods when open-vegetation habitats were available for trans-continental dispersal and at the times when aridification and temperature decline offered favorable circumstances for bee diversification.

## 1. Introduction

Bees constitute a well-established clade of pollen-feeding Hymenoptera presumed to have arisen during the Early to mid-Cretaceous approximately 113-132 Ma (million years ago) (Cardinal and Danforth, 2013). Their origin coincides with the end of the Gondwana breakup and the early diversification of eudicots, the major group of flowering plants (Cardinal and Danforth, 2013). Currently presenting a worldwide distribution and about 20.000 described species (Ascher and Pickering, 2020), bees have been hypothesized to have originated in the Southern Hemisphere, more specifically, in the xeric interior of Gondwana (Michener, 1979; Litman et al., 2011; Hedtke et al., 2013). Over the last 20 years, much progress has been made on our understanding of the origin and evolution of bees mainly due to the use of molecular data and fossil information combined with statistical phylogenetic methods to reconstruct bee phylogenies at different taxonomic levels (see Danforth et al., 2013, for a review). Among the main groups of bees, the Andreninae remain the most poorly understood lineage in terms of phylogenetic relationships of its supraspecific taxa and biogeographic history (Danforth et al., 2013).

In the present work we focus on the phylogenetic and biogeographic history of andrenine bees. This group provides an excellent system for exploring historical biogeographic patterns of biotic interchanges that took place during the Cenozoic mainly between the North and South America regions. Andreninae constitute a well-established clade of bees with about 3,000 species that represent over 15% of all bee diversity (Ascher and Pickering, 2020) and diverged from the clade (Halictinae (Stenotritinae + Colletinae)), around 90 Ma in the late Cretaceous (Cardinal and Danforth, 2013; Sann et al., 2018). The subfamily is widely distributed on all continents, except Australia and tropical Asia, and are exceptionally diverse in the temperate and xeric parts of the Western Hemisphere (Michener, 2007). Among the 61 known genera of andrenine, the New World fauna encompasses around 45 genera, with 31 of them found only in South America. Outside the Americas, Andreninae have a smaller genus-level diversity, with 14 genera reported for the Palearctic region and 5 in the Afrotropics.

Andreninae comprise typically slender medium-sized (6-10 mm long), black with some yellow marks, and sparsely haired bees, but also robust and hairy bees such as Oxaeini (Michener, 2007). It has been consistently recovered as a monophyletic group (Ascher, 2004; Danforth et al., 2004, 2006a, 2006b; Cardinal and Danforth, 2013; Danforth et al., 2013; Hedtke et al., 2013). The most distinctive morphological character is the subantennal area defined by two subantennal sutures, with the presence of facial fovea in females and some males, and the pointed glossa being also useful features in their recognition (Michener, 2007). The subfamily is remarkable for the high proportion of pollen specialists (oligolectic) species, each visiting a few species within the same plant family, such as Asteraceae, Cactaceae, Fabaceae, Malvaceae, Melastomataceae, Onagraceae, Oxalidaceae, Passifloraceae, Solanaceae, and Turneraceae (Michener, 2007). As far as we know, all andrenine species are solitary, however, several groups also show communal nesting biology (such as *Andrena, Oxaea*, Perditini, *Panurgus*, and some Protandrenini) (Michener, 2007). The nests are excavated in the soil with one or a short series of cells at the end of each lateral burrow radiating from the main burrow. Many species are uni- or bivoltine (i.e. producing one or two generations per year), especially in seasonally dry areas, but many common polylectic species are multivoltine. The cleptoparasitic associations of several species of nomadine bees with andrenines are an interesting and complex system to be investigated (Rozen, 1989). In this sense, resolution of the phylogenetic relationship among Andreninae lineages is an important step for understanding the evolution of cleptoparasitism in bees (Rozen, 1992), as well as the evolutionary history of bee-plant associations (Larkin et al., 2008; Danforth et al., 2013).

The classification of Andreninae has a complex history with variable number of recognized internal taxa. For many years the Andreninae was treated as a slightly less-inclusive group, excluding Oxaeini (Ascher et al., 2006; Engel, 2015). Members of this tribe differs considerably from other Andreninae, what lead to its placement as a separate higher taxon based on adult and larval morphology (e.g. Michener, 1944; Rozen, 1964, 1965, 1993; Hurd and Linsley, 1976; Alexander and Michener, 1995). Ruz (1986, 1991) was the first to investigate phylogenetic relationships among genera of the panurgine line, followed by Patiny (1999), who analyzed the relationship among the Old-World members of panurgine based on adult morphological characters proposing six tribes: Panurgini, “Camptopoeumini”, Panurginini, Mermiglossini, Melitturgini, and “Paramelittergini”. The forms included in this arrangement were classified into two tribes (Panurgini and Melitturgini) by Michener (2000, 2007), a proposal that was largely followed by subsequent authors. Later, Ascher (2004) analyzed the whole Andreninae using morphological (adult and larvae) plus EF-1α (F2 copy) sequences, only partially published (in Rozen, 2003) in order to establish the phylogenetic relationship of the enigmatic Nolanomelissini.

Our understanding of the phylogenetic relationships among andrenine bees remains fragmented due to limited taxon sampling mainly from South America – the richest region in diversity of supra specific taxa. Additionally, a global biogeographic analysis based on a well-resolved fossil calibrated phylogeny of the subfamily has not yet been attempted thus far. Thus, the primary goal of the work is to provide a solid phylogenetic framework with special emphasis on the New World, as an important step to establish a stable classification of the Andreninae and to the understanding of their evolution through the Cenozoic. This is the first time the monotypic tribe Protomeliturgini has been included in a molecular phylogenetic work. The current study provides information to understand the ancient diversification that took place in South America and the complex biogeographic scenario and evolutionary factors that have produced the biotic interchange between the warm areas of the Americas during the Mesozoic and Cenozoic.

## 2. Material and methods

### 2.1. Taxon and molecular sampling

The taxon sampling comprises 91 species of the subfamily Andreninae sensu lato, representing all currently recognized suprageneric taxa and the main geographical occurrences within the subfamily. We follow the classification of Melo and Gonçalves (2005) for the main lineages of bees, treated as subfamilies of a single family (Apidae sensu lato), and Moure et al. (2012) for American taxa of Andreninae; in the case of Old-World taxa, we use the subtribes of Ascher and Engel (2017) at tribal level, as originally proposed by Patiny (1999). According to the classification adopted here, Andreninae is subdivided into the following extant tribes: Andrenini, Euherbstiini, Calliopsini, Neffapini, Melitturgini, Mermiglossini, Nolanomelissini, Oxaeini, Panurgini, Panurginini, Perditini, Protandrenini, and Protomeliturgini. The higher-level bee phylogenies indicate a large clade uniting all the short-tongued subfamilies (except Melittinae) (Danforth et al., 2006a, 2006b; Cardinal and Danforth, 2013; Danforth et al., 2013; Hedtke et al., 2013), thus we choose fifteen terminals to represent the subfamilies from this clade: Colletinae (7 spp.), Halictinae (6 spp.) and Stenotritinae (2 spp.). Attention to the choice of species was taken in order to more accurately place fossil calibration points and to maximize morphological diversity and geographic distribution for the ancestral area reconstruction. In total, 106 taxa were included in the analyses, as described in Supplementary Tables S1-S2 along with taxonomy, collection site, voucher information, and GenBank accession numbers (GenBank accession numbers will be indicated upon this manuscript acceptance).

Molecular sequence data were generated for this study fulfilling a huge gap of South American andrenine bees in GenBank. Most newly sequenced specimens were field-collected by us or obtained through donation or loans from other researchers and institutions (see Acknowledgments). In addition, sequence data were obtained from the databases of Genbank and BOLD (Barcode of Life Data System) mostly from Ascher et al. (2001), Ascher (2003, 2004), Danforth et al. (2006a, 2006b), Larkin et al. (2006), Almeida and Danforth (2009), Cardinal and Danforth (2013), and Packer and Ruz (2017). Total DNA was mostly isolated from bees preserved in EtOH, but also from pinned museum specimens collected up to twelve years before processing. Muscular tissue was taken from thorax or legs and the DNA was extracted using the Qiagen DNeasy blood and tissue extraction kit, following the manufacturer’s protocol. In the case of museum species, the whole body was maintained in an insect relaxing chamber for 24-48 hours prior to extraction following the protocol of Evangelista et al. (2017) for museum specimens, using the same extraction kit. Voucher specimens of the newly generated sequences are housed in the DZUP – Entomological Collection “Jesus Santiago Moure” at Federal University of Paraná, Brazil, or at their home institutions (Supplementary Tables S1-S2).

The selected regions comprise four nuclear protein-coding genes: elongation factor-1 F2 copy (EF-1a), long-wavelength rhodopsin (opsin), RNA polymerase (poly) and wingless (wg); two nuclear ribosomal RNA loci: large subunit 28S rRNA (28S) and 18S rRNA (18S); and one mitochondrial protein-coding gene: cytochrome c oxidase subunit I (COI). These genes have provided robust results for several phylogenetic studies of bees at distinct levels of relationships. Newly generated sequences refer only to EF-1a (650 base pairs (bp)), wingless (500 bp), and 28S rRNA (700 bp). Primer information and PCR conditions can be found in electronic supplementary material (Table S3). PCR success was examined by gel electrophoresis on 1% agarose gel and PCR products were purified and sequenced in forward and reverse directions by Macrogen Inc. (South Korea). Sequence assembling was generated with the Staden Package (Staden et al., 2000). Chromatogram quality evaluation and corrections were done in BioEdit v5.0.9 (Hall, 1999).

All genes were separately aligned in MAFFT v. 7 (Katoh et al., 2019) with default parameters: gap opening penalty = 1.53; offset value = 0. The alignment of the protein coding genes EF-1a and opsin relied on the LINS-i strategy, which is recommended for sequences with multiple conserved domains and long gaps. The alignment of the 18S rRNA, wingless and COI relied on the G-INS-i strategy, recommended for sequences with a global homology. The ribosomal 28S rRNA was aligned based on its secondary structure using the Q-INS-i algorithm. Minor adjustments were made by eye in Geneious, and we made sure that the introns/exon boundaries of EF-1a and opsin were maintained. Sequences from the seven gene fragments were concatenated in Sequence Matrix v. 1.7.8 (Vaidya et al., 2010). The final number of aligned nucleotides used in the analyses was 7952 base□pairs in length (EF-1a: 1946 bp; opsin: 1494 bp; wg: 512 bp; poly: 842 pb; 28S rRNA: 1714 bp; 18S rRNA: 786 bp; COI: 658 bp).

### 2.2. Phylogenetic analysis and divergence time estimation

All genes were concatenated into a single matrix subdivided as follows: introns and exons of the protein coding regions EF-1a and opsin, poly, wg, COI, 28S rRNA, and 18S rRNA totalizing 16-character sets. The search for the best partitioning scheme and models of DNA substitution was performed in PartitionFinder v. 2 (Lanfear et al., 2017) based on the Bayesian Information Criterion and the algorithm greedy (see Supplementary Table S4 for specific information of data partition and their best fit models). The phylogenetic tree searches were carried out using the concatenated matrix with nine data partitions under maximum likelihood and Bayesian inferences. Maximum likelihood analysis was performed in RaxML (Stamatakis, 2006) in the CIPRES server (Miller et al., 2011). Simultaneous bootstrap analyses with 1000 replicates were conducted to evaluate node support. Bayesian phylogenetic tree searches were conducted in MrBayes v. 3.2. (Ronquist et al., 2012) in CIPRES. The Markov chain Monte Carlo (MCMC) was run for 20 million generations sampled every 1000^th^ generation. The stationarity of all parameters and convergence of both runs were accessed in Tracer v. 1.7.1 (Rambaut et al., 2018). A 25% burn-in was applied and a 50% majority rule consensus was computed with the remaining trees in TreeAnnotator (part of BEAST 2.0 package, Bouckaert et al., 2014). Resulting ML and Bayesian trees were visualized and edited in FigTree v. 1.4.4 (Rambaut, 2016).

Divergence times were estimated in a Bayesian framework using BEAST 2 (Bouckaert et al., 2014) by employing the same partition□scheme and models of phylogenetic analyses. Four bee fossils were used to calibrate the Andreninae tree, two belonging to this subfamily and the others to Halictinae and Colletinae. Compression fossils attributed to *Andrena* from the Eocene of Florissant, Colorado (Priabonian Age: 37.2–33.9 Ma) were used to calibrate the node of extant *Andrena* species. There are four species from Florissant deposit attributed to *Andrena*: *A. grandipes* Cockerell, 1911; *A. hypolitha* Cockerell, 1908; *A. percontusa* Cockerell, 1914 and *A. sepulta* Cockerell, 1906. Since the descriptions by Cockerell, their identity has not been much investigated, but all have been undoubtedly attributed to the megadiverse genus *Andrena*. For this point, a lognormal distribution prior was applied with offset = 34, stdev = 0.623 and Mean = 1.8, corresponding to a minimum bound of 34 Ma, median value of 40 and 95% quantile of 50. *Heterosarus eickworti* Rozen, 1996 from the Miocene Dominican amber (20.43–13.65 Ma) was used to calibrate the node uniting the two extant species included in our study. The crown group of *Heterosarus* was given a lognormal distribution with offset 14, stdev = 0.432 and M = 1.8, corresponding to a minimum bound of 14 Ma, median value of 20, and 95% quantile of 26.31. Two other species from Dominican amber, *Chilicola (Hylaeosoma) gracilis* Michener and Poinar, 1996 and *Chilicola (Hylaeosoma) electrodominicana* Engel, 1999, were attributed to the crown group of *Chilicola* (Colletinae, Xeromelissini), following the same prior used in *Heterosarus. Electrolictus antiquus* Engel, 2001 from the Eocene Baltic amber was used to calibrate the stem group of Halictini (in Halictinae) given a lognormal distribution prior with the same parameter as the *Andrena* fossils, since the Baltic amber has also a Priabonian age. All geological ages attributed to the fossils mentioned here were retrieved from Fossilworks (Behrensmeyer and Turner, 2013).

A normal prior distribution was attributed to the crown age of Andreninae+(Colletinae, Halictinae) based on previous broader age estimates for Hymenoptera (Aculeata): from Branstetter et al. (2018) (89.3 Ma, 95% HPD: 76.66-100.17 Ma) and Peters et al. (2017) (99, 95% HPD 82-118). An average value between these two estimates was calculated and the calibration point was given a normal distribution, offset 94, sigma 2.5 and mean 1 (5% quantile 90.1, 95% 99.9). The tribes Protandrenini and Mermiglossini were constrained to be monophyletic based on the Maximum Likelihood and Bayesian analyses, which consistently recovered these clades. Supplementary Figure S1 depicts all points of fossil calibration, secondary calibration and monophyletic constraints. Substitution and clock models were unlinked among partitions, but trees were linked. A Yule speciation model was used as tree prior. Markov chain Monte Carlo (MCMC) searches were conducted for 100×10^6^ generations sampled every 10,000 generations with the first 25% discarded as burn-in. Convergence and stationarity of the runs were accessed in Tracer v 1.7.1 (Rambaut et al., 2018) using the ESS scores. The final tree was created in TreeAnnotator (both part of BEAST 2 package). Resulting maximum clade credibility tree was visualized and edited in FigTree v. 1.4.4 (Rambaut, 2016). All trees with associated data matrix will be deposited in TreeBASE upon manuscript acceptance.

### 2.3. Ancestral range estimation

We were interested in illuminating the biogeographic history of Andreninae and identifying their most probable ancestral range and the events of dispersal between the New World and the Old World and between North and South America. For achieving this purpose, we conducted an ancestral range estimation analysis using five biogeographically relevant regions representing the total distribution of Andreninae and outgroups: A. South America, B. North America, C. Palearctic, D. Afrotropical and E. Australian. The Palearctic is here used in broad sense, including some extralimital elements whose distributions extend to the Oriental region. Geographical occurrences that could be assumed to be secondary were considered biogeographically uninformative. For example, the species-rich genus *Andrena* was considered here as occurring only in North America and in the Palearctic region, in spite of known geographic records in the Neotropics, Afrotropics and Oriental region, since its presence in these latter regions has been attributed to younger clades (Pisanty et al., 2021). Biogeographic coding for *Melitturga* considered the known distribution of the whole genus, even though we only sampled palearctic species.

Ancestral ranges of Andreninae were estimated using R package BioGeoBEARS (Matzke, 2013, 2014) comparing different maximum likelihood versions of three classic biogeographic methods: DEC (Dispersal-Extinction-Cladogenesis: Ree and Smith, 2008), BayArea (Bayesian Inference of Historical Biogeography for Discrete Areas: Landis et al., 2013) namely BAYAREALIKE, and DIVA (Dispersal-Vicariance Analysis: Ronquist, 1997) namely DIVALIKE. The maximum clade credibility tree derived from the BEAST analysis was used as input and each terminal taxon was coded for presence/absence in the five geographic areas (Supplementary Table S5). We applied a dispersal multipliers matrix to give different probabilities for different events of dispersal between continents. Dispersals between, for example, South America and Africa, which is very implausible for Andreninae bees after Gondwana breakup, were given a 0 probability; while reasonable events, for example well-documented long-distance dispersals between South and North America, before the closure of the Panama Isthmus, were given a 1 probability. We assessed the overall fitness of the models with likelihood-ratio tests and AIC values.

## 3. Results

### 3.1. Phylogenetic Relationships and Divergence Time Estimation

We assembled the largest molecular dataset for andrenine bees up to date: our final matrix contains 7952 nucleotides and 106 taxa, comprising approximately 70% of the currently recognized genera in Andreninae *sensu lato*. The monophyly of most tribes and many of their relationships are well-supported based on bootstrap proportions and Bayesian posterior probabilities (i.e. node supports higher than ≥ 70% ML bootstrap and ≥ 98% Bayesian posterior probability). Figure 1 and Supplementary Figure S2 depict the phylogeny of Andreninae recovered by ML and Bayesian inference, respectively. Figure 2 and Supplementary Figure S3 show the fossil-calibrated tree and ages for main nodes and related 95% highest posterior probability intervals are listed in Table 1.

**Figure 1.**
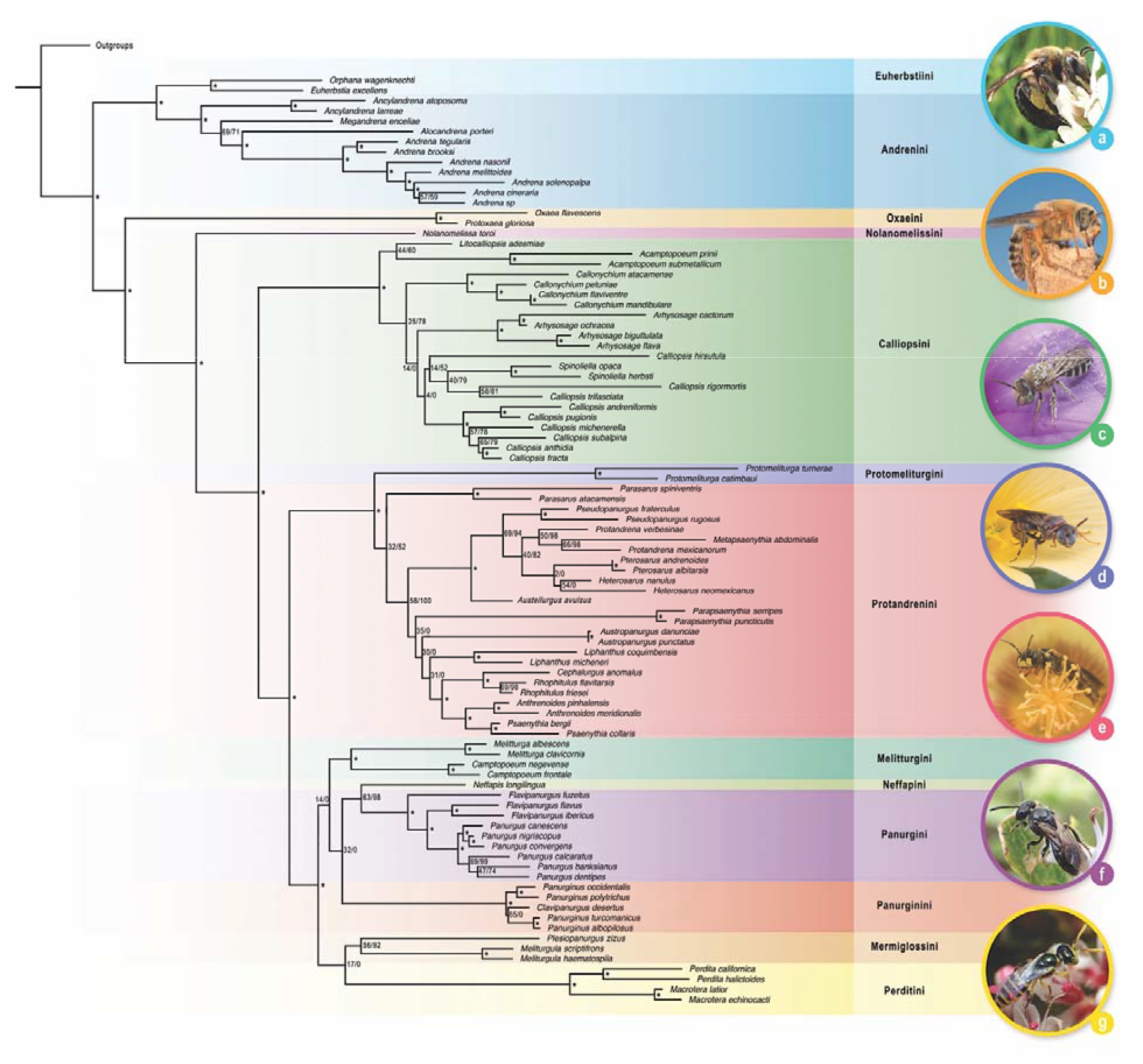
Phylogenetic relationships among 91 species of Andreninae bees resulting from a Maximum Likelihood analysis of concatenated data set of the following seven gene loci: elongation factor-1a F2 copy, long wavelength rhodopsin, wingless, RNA polymerase II, 28S rRNA, 18S rRNA, and cytochrome c oxidase subunit I. Complete tree of Maximum likelihood and Bayesian analysis are shown as Figure S1 and S2, respectively. Maximum likelihood bootstrap support (BS) and Bayesian posterior probability (BPP) values are shown at nodes. Clade support ≥98% BPP and ≥70% BS is indicated by asterisk on subtending node. (a) *Andrena vicina*, female. (b) *Oxaea* sp., male. (c) *Acamptopoeum prinii*, male. (d) *Protomeliturga catimbaui*, female. (e) *Cephalurgus anomalus*, male. (f) *Panurginus* sp., female. (g) *Perdita rhois*, male. Photo credits: Tom Barnes (a, f, g); Julio Pupim and Adriana Tiba (b, c, e); Clemens Schlindwein (d).

**Figure 2.**
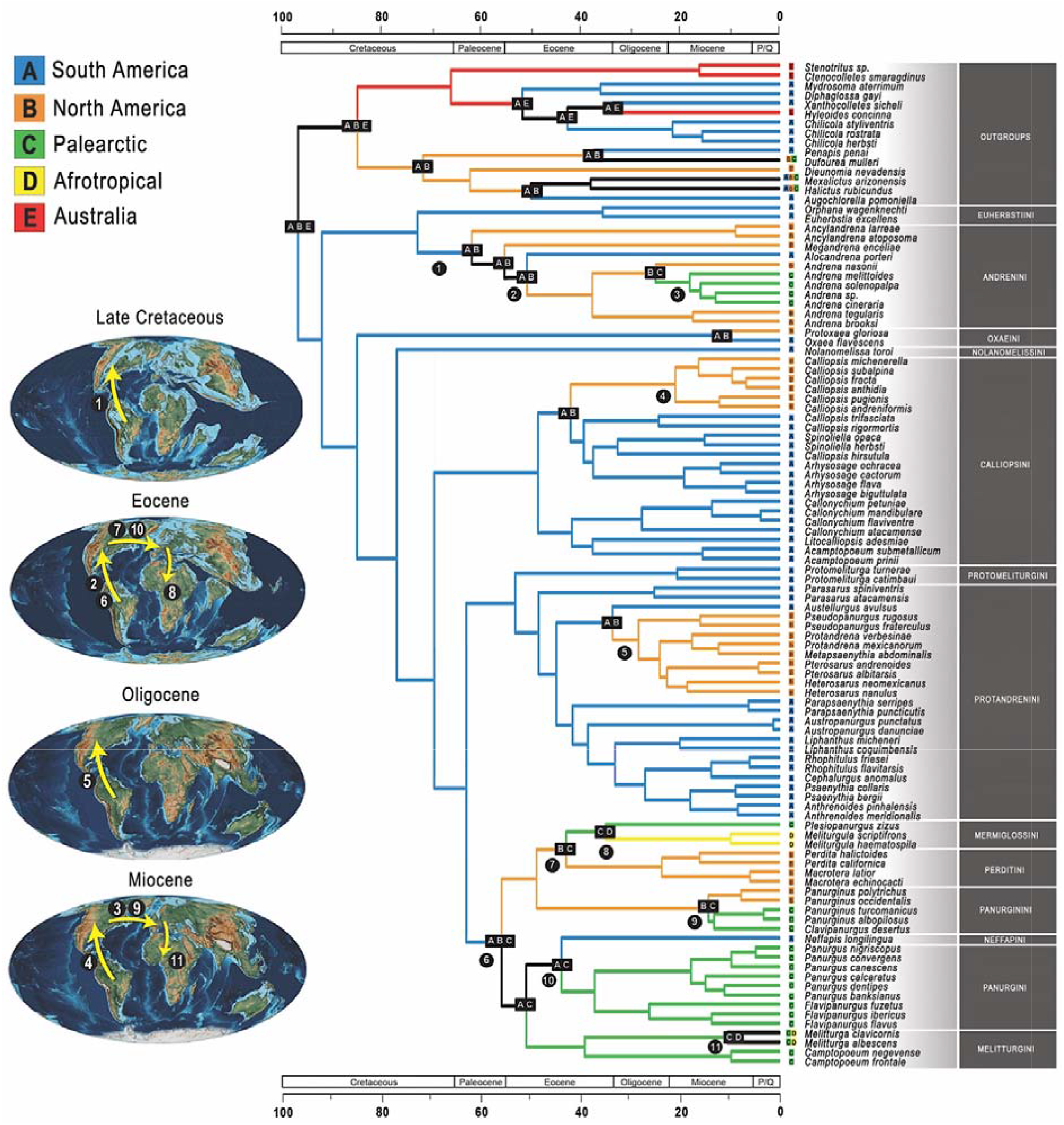
Maximum likelihood ancestral range estimation in Andreninae, using the best model DEC (dispersal-extinction-cladogenesis model) implemented in BioGeoBEARS. The pie diagrams at nodes show the relative probability of the possible states (areas or combinations of areas). Color boxes show the distribution of each taxa within the clade. Black boxes represent nodes at which more than an ancestral area was recovered. Black circles represent nodes at which lineage-interchange events may have occurred. Maps adapted from Scotese (2014).

**Table 1.** Divergence time estimates for crown and stem groups of most relevant clades of Andreninae bees. Values in parentheses refers to 95% Highest Posterior Density intervals. Ages in million years (My).

| Clade | Crown age | Stem age |
| --- | --- | --- |
| Andreninae | 91 (82–99) | 96 (91–101) |
| Andrenini + Euherbstiini | 72 (59–86) | 91 (82–99) |
| Andrenini | 61 (51–74) | 72 (59–86) |
| *Alocandrena* + *Andrena* | 50 (40–58) | 55 (45–65) |
| Euherbstiini | 35 (11–61) | 72 (59–86) |
| Oxaeini | 11 (2–23) | 84 (75–94) |
| Nolanomelissini | - | 76 (67–86) |
| Calliopsini | 48 (37–59) | 69 (59–79) |
| *Calliopsis* (North America) | 20 (12–31) | 42 (32–53) |
| Protomeliturgini + Protandrenini | 53 (43–62) | 63 (53–72) |
| Protomeliturgini | 20 (8–34) | 53 (43–62) |
| Protandrenini | 48 (39–57) | 53 (43–62) |
| Protandrenini (North America) | 28 (22–34) | 33 (25–41) |
| Holarctic-Afrotropical clade* | 56 (46–65) | 63 (53–72) |
| Melitturgini | 39 (26–51) | 50 (38–58) |
| Mermiglossini | 34 (21–47) | 42 (32–52) |
| Neffapini | - | 43 (28–50) |
| Neffapini + Panurgini | 43 (28–50) | 50 (38–58) |
| Panurgini | 37 (19–40) | 43 (28–50) |
| Panurginini | 14 (7–21) | 48 (38–59) |
| Perditini | 23 (14–32) | 42 (32–52) |
| * It includes also the Chilean tribe Neffapini; the whole group could be referred as the supertribe Panurgidi Leach |  |  |

All analyses recovered the monophyly of Andreninae sensu lato with high support. The divergence time estimation indicates that Andreninae diverged from the clade Halictinae (Colletinae, Stenotritinae) in the Late Cretaceous and evolved rapidly soon afterwards, considering the age of 96 (91–101) Ma for the crown group. Most andrenine tribes were recovered as monophyletic with maximum support, except for tribes Melitturgini (99 BPP, 72% BS) and Panurgini (100 BPP, 88% BS) and the not well supported Protandrenini and Mermiglossini. Furthermore, the position of all tribes agrees between ML and BI, expect for Panurginini and Mermiglossini.

The basalmost divergence within Andreninae involves two major clades, both containing widespread species-rich subclades. The first split corresponds to a clade formed by the mainly Holarctic species-rich tribe Andrenini and an endemic Chilean group, the Euherbstiini originated around 73 (59–86) Ma, in the Late Cretaceous. The monophyly of Andrenini is recovered encompassing the monospecific genus *Alocandrena*, endemic to Peru, as sister group of the speciose *Andrena*, a genus widely distributed in the Holarctic region. The second major clade in Andreninae contains a series of tribes, involving both New- and Old-World elements. The base of this clade contains a grade with two New World tribes, the Oxaeini and the Nolanomelissini, which have diverged from the remaining tribes quite early in the Late Cretaceous. Oxaeini is an ancient lineage, having diverged in the Campanian period (∼85 Ma), but whose crown is estimated to be much younger. Nolanomelissini, a monospecific group from Chile, is very distinctive and diverged from the panurgine line (see below) around 77 (67–87) Ma.

The remaining tribes of this second major clade form a group treated as the Panurginae in Michener’s classification and that we refer here as the panurgine line. This large assemblage contains Calliopsini, splitting at the base as sister group of the large clade formed by Protandrenini, Protomeliturgini, Perditini, Melitturgini, Mermiglossini, Neffapini, Panurginini and Panurgini. The divergence between Calliopsini and the other tribes of the panurgine line is estimated to have occurred at 69 (60–79) Ma, while the crown age of Calliopsini is 20 million years younger (Fig. 2 and Table 1). The relictual tribe Protomeliturgini, endemic to the dry areas of northeastern Brazil, is recovered with strong support as sister clade of Protandrenini. The monophyly of Protandrenini is confirmed, although with low support, excluding the monotypic genus *Neffapis* which proved to be phylogenetically related to the Panurgini. Within Protandrenini, the North American genera *Heterosarus, Protandrena, Pseudopanurgus* and *Pterosarus* were shown to form a clade that diverged from de South American genera in the early Oligocene, around 33 (26–41) Ma (Fig. 2).

The remaining tribes of the panurgine line form a large clade distributed in the Afrotropical (Mermiglossini), Nearctic (Perditini and Panurginini), and Palearctic (Panurgini, Panurginini and Melitturgini) regions, except for Neffapini, a monotypic tribe known only from northern Chile. Phylogenetic relationships among these tribes remain somewhat uncertain, varying to some degree depending on the method used to analyze the data, probably due to the short length of the involved branches (Fig. 1 and Supplementary Fig. S1). The sister-group relationship between Neffapini and Panurgini is recovered in all analyses (79% BS, 100 BPP) and their crown age estimates correspond to about 44 (28–50) Ma.

Our analyses recovered the monophyly of all non-monotypic genera, except *Calliopsis* (Calliopsini) and *Heterosarus* (Protandrenini). This last one was placed in a trichotomy with *Pterosarus* by the Bayesian analysis, but monophyletic in the ML results. *Calliopsis* is paraphyletic, with the species endemic to South America forming a clade distantly related to those from the Nearctic region. Most of the genera in Andreninae (∼65%) had their origin in the Miocene (Fig. 2).

#### Ancestral Range Estimation

The model DEC yielded the highest likelihood and best AICc scores for ancestral range estimation for the Andreninae phylogeny. Statistical results for all three applied models are given in Supplementary Table S6. Ancestral range estimates plotted in the Maximum Clade Credibility tree are show in Figure 2, for the most important nodes, and in Supplementary Figure S3 for all nodes. The reconstructed scenario shows unequivocally that the most recent common ancestor (MRCA) of the subfamily Andreninae originated in South America in the Late Cretaceous. The early evolution of its main lineages took place in this continent until the end of the Cretaceous, when the first event of dispersal to North America occurred in the Maastrichtian involving the lineage that gave rise to the tribe Andrenini. Other incursions from South to North America and vice-versa involved different lineages in the Eocene within Andrenini and the panurgine line, and during the Oligocene within Protandrenini and Calliopsini, with the last two tribes retaining a largely South American distribution. This indicates at least three periods of long-distance dispersal or possible land connections between the two continents, much prior to the closure of the Panama Isthmus.

## 4. Discussion

### 4.1. Phylogeny and Higher Classification of Andrenine Bees

Andreninae is recovered as a well-supported monophyletic group, including the Oxaeini, as previously proposed (Ascher, 2003, 2004; Danforth et al., 2006a, 2006b; Cardinal and Danforth, 2013; Hedtke et al., 2013). One of the earliest-branching lineages of Andreninae corresponds to the already previously recognized clade uniting the tribes Andrenini and Euherbstiini (Pisanty et al., 2021). Our results show *Alocandrena*, a monotypic genus from the Peruvian Andes, nested within Andrenini. Due to its unusual morphology in relation to the other Andrenini, Michener (2000) placed *Alocandrena* in a separate higher taxon, a classificatory decision corroborated neither by the present study nor by previous molecular phylogenies (Pisanty et al., 2021). The phylogenetic relationships among the genera of Andrenini remain controversial. We found *Ancylandrena* (*Megandrena* (*Alocandrena, Andrena*)), while Pisanty et al. (2021) recovered *Andrena* as sister to *Alocandrena* (*Ancylandrena, Megandrena*).

In the other basalmost clade of Andreninae, we have Oxaeini recovered as sister to the remainder of this clade. This tribe is composed of 22 species of quite large and fast hover flying bees restricted to warmer regions of the Western Hemisphere (Ascher et al., 2006; Engel, 2015). The tribe is considerably different morphologically from the other Andreninae, what lead to its placement as a separated family of the traditional classification (e.g. Rozen, 1964, 1965, 1993; Hurd and Linsley, 1976), mainly on the basis of several distinct features of their mature larvae and adults (Hurd and Linsley, 1976; Michener, 2007; Engel, 2015). However, broader studies using both larval and adult morphological characters, strongly support the position found here with the group nested within Andreninae sensu lato (Michener, 1944; Graf, 1966; Rozen, 1993, 1994; Alexander and Michener, 1995; Ascher, 2003, 2004; Danforth et al., 2006a, 2006b; Hedtke et al., 2013; Engel, 2015).

The enigmatic species *Nolanomelissa toroi*, known only from the southern border of the Atacama Desert in Chile, appears as sister to remaining tribes of the panurgine line. This species is oligolectic on pollen of *Nolana rostrata* (Solanaceae) (Rozen, 2003; Michener, 2007), and has a number of morphological apomorphies, but also shares characteristics with different tribes of Andreninae making it difficult to assign to a tribe unequivocally (Rozen, 2003). However, our results agree with the placement of *Nolanomelissa* in its own tribe, Nolanomelissini, as proposed by Rozen and Ascher (in Rozen, 2003). Calliopsini, here represented by all of its genera (except *Xeranthrena*) and including South American species of *Calliopsis* is recovered as monophyletic. Phylogenetic relationships based in adult morphology have recovered a clade containing the genera *Callonychium, Spinoliella, Xeranthrena* and *Arhysosage* excluding *Calliopsis, Acamptopoeum* and *Litocalliopsis* (Ruz, 1991; Roig-Alsina and Compagnucci, 2003; Gonzalez et al., 2017). Data on larval morphology have also supported a close relationship between *Arhysosage, Callonychium* and *Spinoliella* (Rozen, 2013; referred by him as the *Spinoliella* group). Our results, however, indicate that *Calliopsis* sensu lato belongs to this group and *Spinoliella* makes the large genus *Calliopsis* paraphyletic.

Species of *Calliopsis* are classified in nine subgenera, three of them endemic to South America and six to North America. Although Ruz (1991) evidenced the monophyly of *Calliopsis* sensu lato, she highlighted the South American *C*. (*Liopoeum*) as one of the most distinctive subgenera, with some species having the female metasomal terga with hair bands and others with pigmented yellow bands. It seems clear that the genus *Calliopsis* still require a more thorough phylogenetic investigation, with a broader taxonomic sampling to understand the limits and definitions of the endemic South American subgenera in order to define their taxonomic status. *Litocalliopsis* was corroborated as a distinct lineage sister to *Acamptopoeum*, contradicting previous hypothesis proposing it as sister to *Calliopsis* (Roig-Alsina and Compagnucci, 2003). In fact, these authors have also indicated several morphological features shared between *Litocalliopsis* and *Acamptopoeum*.

Our molecular phylogeny provides a novel sister-group relationship between Protomeliturgini and Protandrenini with maximum support. Previous hypotheses suggested Protomeliturgini as close to Perditini and Calliopsini (Ruz, 1986, 1991). Protomeliturgini contains a single genus with two described species distributed in semiarid areas of northeastern Brazil and oligolectic in flowers of Turneraceae (Medeiros and Schlindwein, 2003). *Protomeliturga* is easily distinguished from other lineages by the very elongated first two articles of the labial palpus, strongly curved 2^nd^ abscissa of the vein M (basal vein), and tergum 7 of male strongly curved forward, with a pair of apicolateral teeth (Schlindwein and Moure, 2005; Michener, 2007).

Protandrenini is morphologically heterogenous, therefore not obviously monophyletic, but was recovered by the present molecular dataset. Our results unequivocally exclude *Neffapis* from Protandrenini. The placement of *Austellurgus avulsus* (Ramos & Melo, 2006) as sister to the North American Protandrenini is here reinforced. Michener (2007) gave *Protandrena* an overly broad scope, recognizing as subgenera the South American species of *Austropanurgus* and *Parasarus* and North American species of *Heterosarus, Metapsaenythia*, and *Pterosarus*. Yet, *Pseudopanurgus* was also previously recognized in a much broader sense synonymizing the genera *Heterosarus, Pterosarus*, and *Xenopanurgus*, but based on an incomplete sampling of those lineages. Our results do not provide support to these propositions, showing *Protandrena* sensu Michener (2007) as polyphyletic with respect to genera from South America and *Pseudopanurgus* sensu Ascher (2004) paraphyletic in relation to *Protandrena*. We have also found out *Protandrena* sensu stricto paraphyletic to *Metapsaenythia* as already indicated by both morphological and molecular data (Ascher, 2004; Michener, 2007).

Within the panurgine line, we also found a large clade formed by the tribes Neffapini, Perditini, Mermiglossini, Melitturgini, Panurgini and Panurginini (see below for decision on classification system). Relationships within this large and heterogeneous clade, containing both New and Old World taxa, differed somewhat between the analyses, with the alternative arrangements involving mainly Panurginini and Mermiglossini. The monospecific Neffapini, the only South American element in this clade, came out as sister-group to the Old World *Flavipanurgus* and *Panurgus* (Panurgini). *Neffapis*, endemic to Coquimban desert in Chile, exhibits many unique characters such as the extremely long glossa and third labial palpus and a minute maxillary palpus with only two palpomeres (Rozen and Ruz, 1995), which led to difficulties in positioning it within existing tribes. Proposal of a separate tribe for *Neffapis* by Ascher (in Engel, 2005) is corroborated here, although in a rather different phylogenetic scenario.

Monophyly of Perditini and its two genera, *Perdita* and *Macrotera*, corroborates morphological phylogenies (Danforth, 1996) and recognition of *Macrotera* as a valid genus distinct from *Perdita*. This tribe is close to Palearctic and Afrotropical tribes (Patiny, 1999; Ascher, 2003). The phylogenetic relationships of *Camptopoeum* with *Melitturga* and *Meliturgula* with *Plesiopanurgus* corroborate the establishment of the tribes Melitturgini and Mermiglossini, respectively. It is possible that this large clade may also include the intriguing species *Simpanurgus phyllopodus* (Warncke, 1972), endemic to the Iberian Peninsula and known only from males. Previous treatments indicated *Simpanurgus* as a subgenus of *Panurgus* (Michener, 2007) or as *incertae sedis* in relation to the Old-World tribes (Ascher and Engel, 2017). The inclusion of molecular data from this species, as well as a broader sampling of Old-World elements, would improve our understanding of its position in the phylogeny of Andreninae.

The classification system adopted here for this large clade of the panurgine line gives tribal status to the main lineages. The systematic position of the genera within this large clade varies in different papers and differs according to the suprageneric-level classification adopted (Table 2). A comprehensive background about the distinct classification systems adopted for these taxa is provided by Ascher and Engel (2017). Here we propose the following taxonomic arrangements: Mermiglossini include *Plesiopanurgus* and *Meliturgula*, represented here by a total of three species, and the non-represented genera *Flavomeliturgula, Gasparinahla* and *Mermiglossa*; Melitturgini include *Camptopoeum, Melitturga* and the genera not sampled here *Avpanurgus* and *Borgatomelissa*; Panurgini include *Panurgus* and *Flavipanurgus*; Panurginini include only *Panurginus*; and Perditini comprise *Macrotera* and *Perdita*. We disagree from previous classifications that give status of subtribe to these lineages, as proposed by Ascher (2004), Engel (2005), and Ascher and Engel (2017), based on the following reasons: (1) We found no evidence for a monoplyletic Panurgini sensu lato; Neffapini would have to be included to make it monophyletic; (2) Use of tribal level preserves the status of well-known groups, as the Perditini; (3) There is a large amount of heterogeneity within the clade, surpassing that found in other andrenine lineages, to assemble them under a single tribe; (4) The first divergences within the clade, dated from the early Eocene, have ages comparable to those of other lineages given tribal status within the Andreninae. The classification system adopted here is summarized in Table 2.

**Table 2.** A comparison of suprageneric-level treatments for Andreninae classification.

| Patiny (1999) | Ascher (2004) | Michener (2007) | Ascher and Engel (2017) | Present study |
| --- | --- | --- | --- | --- |
| NT | Andrenini | Andreninae<sup>1</sup> | NT | Andrenini |
| NT | Euherbstiini | Andreninae | NT | Euherbstiini |
| NT | Oxaeinae | Oxaeinae | NT | Oxaeini |
| then undescribed | Nolanomelissini | Nolanomelissini | Nolanomelissini | Nolanomelissini |
| Calliopsini | Calliopsini | Calliopsini | Calliopsini | Calliopsini |
| Protandrenini | Protandrenini | Protandrenini | Protandrenini | Protandrenini |
| NT | Panurgini: Protomeliturgina | Protomeliturgini | Protomeliturgini | Protomeliturgini |
| NT | Panurgini: Neffapina | Protandrenini | Neffapini | Neffapini |
| Camptopoeumini*<sup>2</sup>,<br>Melitturgini* | Panurgini:<br>Camptopoeumina*<sup>2</sup>,<br>Melitturgina* | Panurgini*,<br>Melitturgini* | Panurgini: Camptopoeina*,<br>Melitturgina* | Melitturgini |
| Mermiglossini*,<br>Paramelitturgini*<sup>3</sup> | Panurgini: Mermiglossina*,<br>Meliturgulina* | Melitturgini | Panurgini: Mermiglossina*,<br>Meliturgulina* | Mermiglossini |
| Panurgini | Panurgini: Panurgina | Panurgini | Panurgini: Panurgina | Panurgini |
| Panurginini | Panurgini: Panurginina | Panurgini | Panurgini: Panurginina | Panurginini |
| Perditini | Panurgini: Perditina | Perditini | Panurgini: Perditina | Perditini |

NT: not treated. \* In part. <sup>1</sup>*Alocandrena* excluded. <sup>2</sup>*Nomen imperfectum* (Engel 2005).
<sup>3</sup>Invalid name (Engel, 2001)

### 4.2. Early evolution of the Andreninae

Our fossil calibrated tree and ancestral area estimation indicate that Andreninae arose in South America during the Turonian, around 90 Ma in the Late Cretaceous, an age consistent with previous estimates (Cardinal and Danforth, 2013; Sann et al., 2018). The biogeographic history of Andreninae involved multiple northward transcontinental dispersal events, from South to North America, with subsequent incursions to the Palearctic region and from there to the Afrotropics, as a result of lineage interchanges between these landmasses. Our results, therefore, suggest that the breakup of Gondwana seem to have had minor impact on the early evolution of Andreninae.

Distribution, relationships, and divergence times among andrenine tribes suggest a pattern of diversification mainly related to historical connections between North and South America. The andrenine tribes originated during a relatively long time-window between the Maastrichtian in the Late Cretaceous (crown age of Andrenini: 72 Ma) and the Miocene (crown age of Oxaeini: 11 Ma). These exchanges between the land masses of the Western Hemisphere gave rise to what is known as the amphitropical distributional pattern (see Michener, 1979), which is exhibited by several lineages of Andreninae, as well as many other New World bee groups.

Aside from land connections, dispersal of andrenine bees across Mesoamerica would have required the availability of suitable habitats. Here we were able to reconstruct a more specific biogeographic scenario showing the early differentiation of Andreninae taking place under conditions similar to those prevalent today in the xeric regions of western South America. Although a large diversity is currently distributed in open vegetation of the American continent, arid and semiarid regions with Mediterranean climate played an important role in the diversification of andrenine bees, especially the western portion of South America (Chile and Argentina) and southwestern North America (Californian deserts and the desertic regions of the southwestern United States and northern Mexico). The importance of these areas to the evolution of Andreninae is supported by the occurrence of many endemic taxa (Michener, 1979; Simpson and Neff, 1985; Turchetto-Zolet et al., 2013). Occurrence of the ancient relictual lineages Euherbstiini, Nolanomelissini and Neffapini west of the Andes in the southern Atacama and Coquimban deserts of Chile is a further evidence of these areas as refugia for bees. The complex geological scenario of Andean orogeny, a long process beginning in the Paleogene, has affected the climate and biodiversity in South America (Hoorn et al., 2010), favoring a progressive formation of xeric habits along western portions of South America (Armijo et al., 2015) and acting as a barrier to bee dispersal. This picture of Andean formation is consistent with our data that shows that the first mountain chain may have acted as isolation barrier for earlier andrenine lineages as Euherbstiini, *Nolanomelissa, Neffapis*, and *Alocandrena*, and also as a strong barrier that prevented the spread of eastern taxa such as the Oxaeini.

### 4.3. Bee faunal interchanges between South and North America

The evolutionary history of the Andreninae involves multiple faunal interchanges between the South and North Americas in at least three different periods suggesting the existence of land connections prior to the final closure of the Panama Isthmus in the Miocene. The time range of the events estimated in our dating analyses is chronologically consistent with different hypotheses of the geological history of connections between the two large landmasses of the American continent from the Late Cretaceous onward. Contrary to Michener’s (1979) hypotheses, we found evidence that south-to-north dispersal of bees is older than the reverse direction.

The early divergence between the tribes Andrenini and Euherbstiini in the Late Cretaceous involved a geodispersal to North America (Fig. 2, Clade 1). Examples of similar exchanges during this period have been reconstructed in other insect groups, such as riodinid butterflies (Espeland et al., 2015), melanopline grasshoppers (Chintauan-Marquier et al., 2011), and in bees of both eucerine and apine lines (see Martins and Melo, 2016). This exchange event between the Americas, during the Campanian and the Maastrichtian, has been named as the First American Biotic Interchange (FABI), and has been originally proposed from patterns found in vertebrates (Goin et al., 2012).

Distinct relative periods of isolation and contact of these landmasses from the Cretaceous to the Pleistocene, and its impact on the amphitropical fauna and flora diversity have long been a subject of discussion. In this way, the disjunct distribution in the American continent implies that Central America served as an important setting for diversification of such taxa in the past. Historical connections between North and South America involve a complex geodynamic of the Caribbean plate that modulated the relative isolation of such areas during distinct geological periods (Iturralde-Vinent and MacPhee, 1999; Ortiz-Jaureguizar and Pascual, 2007; Pindell and Kennan, 2009; Woodburne, 2010; Cody et al., 2010; Farris et al., 2011; Giunta and Orioli, 2011; Coates and Stallard, 2013; Bacon et al., 2013, 2015; Montes et al., 2012, 2015; Cione et al., 2015). Since there is no evidence for an arid corridor through Central America or a full connection of the North and South landmasses before the closure of the Panama Isthmus, geodispersal events should be invoked to explain the diversification of the andrenine bees in the Americas, which perhaps have been facilitated by transient arid conditions in the Central America terrains.

The sister relationships between the Peruvian genus *Alocandrena* and the Holarctic *Andrena* represent another trans-Caribbean geodispersal event, but estimated here to have occurred in the Eocene (∼50 Ma) (Fig. 2, Clade 2). Pisanty et al. (2021) also recovered an early Eocene age for the split between *Alocandrena* and their North American relatives and explained this pattern by long distance wind dispersal through the Central American Seaway, possibly aided by island hopping. Our results also indicate another ancient dispersal event taking place at the beginning of the Eocene and involving the South American lineage that gave rise to a large clade presently distributed in the Nearctic, Palearctic and Afrotropical regions, but also containing a single South American genus, the Chilean *Neffapis*, nested within it (Fig. 2, Clade 6). The distribution pattern exhibited by this clade implies biogeographic scenarios invoking a direct dispersal from South America to the Palearctic region. Such direct connections are unlike to have occurred during this timeframe and considering the presence of some elements in the Nearctic region, as well the Chilean lineage, we presuppose that this ancestral lineage also entered the Old World following a route through North America. This South America–Palearctic disjunct scenario, however, presupposes some extinctions in the Nearctic region since only a few lineages of this clade have North American representatives. Inter-American exchanges during the same time period in the early Eocene have previously been documented for the bee tribes Halictini and Sphecodini (Danforth et al., 2004).

Additional inter-American exchanges occurring under younger ages have been also recovered in our study. Within the tribes Calliopsini and Protandrenini, we found a repeated biogeographic pattern with incursions to North America occurring between 42–28 Ma (late Eocene to early Oligocene) and 33–20 Ma (early Oligocene to early Miocene), respectively (Fig. 2, Clades 4 and 5). The tribe Oxaeini also exhibits an amphitropical distribution and likely represents another dispersal event within this scenario (Fig. 2), but whose divergence estimation needs further investigation with a more comprehensive taxon sampling.

The majority of the most recent and well-documented episodes of multitaxon interchanges between North and South America occurred after formation of a permanent land corridor in the Pliocene (∼3 Ma). This biogeographic event – the Great American Biotic Interchange (GABI) – resulted from the rise of the Isthmus of Panama and climatic changes (mainly glaciations) that allowed many animal groups to cross the Panamanian connection (Simpson and Neff, 1985; Cody et al., 2010; Goin et al., 2012; Wilson et al., 2014; Cione et al., 2015). On the other hand, our results support that andrenine bees have migrated during Oligocene and Miocene along Central America around times when patches of open habitats might have been available. Similar to the Andreninae, species in the bee genera *Diadasia* and *Centris* are also associated with arid regions and their evolutionary history endorses a scenario with prevailing climatic conditions and plant formations adapted to dry conditions in the connecting areas between the American landmasses (Wilson et al., 2014; Martins and Melo, 2016). Hines (2008) also suggested that bumblebees may have arrived in South America in the Miocene crossing Central America when favorable patches of temperate habitats arose.

The GABI was initially characterized by the movement of land mammals between North and South America which requires a contiguous area for overland migration (reviewed by Goin et al., 2012, 2015). Several studies suggest that inter-American connections and interchange of organisms, after Eocene age, could have happened earlier than the final closure of the Isthmus of Panama. Cody et al. (2010) reveal the asynchrony between the evolutionary history of several animal taxa related to GABI and plant migrations from South to North America (at least 20 Ma earlier). As observed for plants, northward transcontinental dispersal events around this same time window have been reconstructed here for the Calliopsini and the Protandrenini, and have also been found in other bee groups [see Ramírez et al. (2010) for early Miocene divergences within the orchid-bee genera *Euglossa* and *Eufriesea*, Almeida et al. (2012) for the colletine *Caupolicana* and *Eulonchopria*, Wilson et al. (2014) for the emphorine genus *Diadasia* and Martins and Melo (2016) for the oil-bee genus *Centris*].

Geological favorable scenarios invoked to understand the biotic movements through the American continent around Oligocene-Miocene include overwater long-distance dispersal (Michener, 1979; Cody et al., 2010), volcanic island hopping (Iturralde-Vinent and MacPhee, 1999; Sturge et al., 2009) or through emergent peninsulas formed in Central America (Kirby et al., 2008; Monte et al., 2012, 2015). Although there is still little evidence for an earlier age for the closure of the Isthmus of Panama sensu stricto as highlighted by O’Dea et al. (2016), the current biological evidence, including that brought in the present contribution, endorse a pattern of early interchanges across Central America much before the closure of the Isthmus of Panama has been completed.

### 4.4. Into the Palearctic and Afrotropical regions

The evolutionary history of the Andreninae occurring in the Old World is explained by our data through four dispersal events from North America to the Palearctic region in two distinct historical periods. Most ancient dispersal movements occurred in the Eocene and involve the evolutionary history of Mermiglossini, which seems to have derived from a North American ancestor common to Perditini, and also of the lineage that gave rise to the clade containing Melitturgini, Panurgini and Neffapini. In both cases, the incursions to the Palearctic region imply a North American MRCA. We also find evidence of a biotic connection between North America and the Palearctic region during the late Oligocene (∼24 Ma) and early Miocene (∼14 Ma) indicated by diversification episodes within of the Holarctic genera *Andrena* and *Panurginus* (Fig. 2, Clades 3 and 8, respectively). The presence of Panurgini and Panurginini in Eurasia via North America was also early inferred by Michener (1979) and Ruz (1986).

Evidence for these two dispersal pulses among the Northern Hemisphere landmasses has been previously advocated by Praz and Packer (2014) for the long-tongued bee tribes Ancylaini and Eucerini. These authors reconstructed a first geodispersal event involving the origin of the clade composed by the genera *Ancyla* and *Tarsalia* reaching the Old World from the New World during the early Eocene, and a second distinct period of exchanges between the Eastern and Western Hemispheres in the Miocene involving some eucerine genera. Intercontinental dispersal events in the opposite direction, i.e. from the Palearctics to the Nearctics, during Miocene age is also inferred to explain the evolutionary history of bumblebees (Hines 2008) and honeybees (Engel et al., 2009).

Expansion of New World terrestrial biotic elements onto the Old World, through a North American route, suggests the existence of Northern Hemisphere land bridges with suitable environmental conditions in distinct ages, such as the Beringia, Thulean and De Geer land bridges (Sanmartín et al., 2001; Brikiatis, 2014). The Bering route functioned in warm periods connecting East Asia and North America. The De Geer Bridge chronologically coincides with the Bering route but connected Eastern North America to Eurasia through the Greenland to Fennoscandia (Sanmartín et al., 2001). The Thulean Bridge was also an intercontinental land bridge that connected the Northern Hemisphere via Greenland but became established well after the interruption of the De Geer route, offering a southerly connection through Greenland, Iceland, Faroe Islands and Scotland. In addition to land connections, the progressive aridification and temperature decline documented during the middle Eocene to early Oligocene led to favorable circumstances for bee diversification. This particular climatic condition that caused the displacement of tropical and subtropical forests led also to further expansion of temperate savanna-like vegetation to as far north as the Beringian and North Atlantic Land bridges in North America (Graham, 1999; Cardinal, 2018).

As similarly inferred by Praz and Packer (2014) for the tribe Ancylaini, the incursion of andrenine bees into the Old Word during the Eocene possibly occurred through the Thulean route, taking into account that it implies warmer environmental conditions, due its more southern position, and its connections to the western Palaearctics where the andrenine bees are more diverse today. Additional examples of geodispersals related to the Thulean route in the same period and invoking similar environmental conditions are known from other groups of organisms (see Praz and Packer, 2014).

Afrotropical distribution of *Melitturga* (Melitturgini) and *Meliturgula* (Mermiglossini) are estimated by our analyses as deriving from two distinct events involving Palearctic ancestral groups. While the event involving the Melitturgini seems to have taken place during the Tortonian age in the Miocene (around 10 Ma; 3-18 HPD 95%), that in the Mermiglossini occurred at a much older age (Fig. 2, Clade 8). Indeed, the scenario within the Mermiglossini is more complex and involves the Afrotropical genus *Mermiglossa*, not sampled here, known from Namibia to central East Africa (Ascher and Engel, 2017). *Mermiglossa* is closely related to *Plesiopanurgus* based on morphological data (Ruz, 1986; Patiny, 1999; Ascher, 2004; Michener, 2007; Ascher and Engel, 2017) what indicates that its presence in the Afrotropics derives from an ancient occupation of Palearctic arid areas in northern Africa and the Middle East by the ancestral lineage of the Mermiglossini.

## 5. Conclusions

In the current study we present the first most comprehensive dated phylogeny to understanding of the complex evolutionary history of Andreninae bees. We provide molecular phylogenetic evidence that corroborate the monophyly of several tribes currently recognized based on morphological evidence. The relationships within the large clade containing the tribes Neffapini, Melitturgini, Mermiglossini, Perditini, Panurgini and Panurginini remain poorly resolved and will require further investigation. The reconstructed evolutionary scenario for the Andreninae sensu lato also provided new insights into the biogeographic patterns and revealed repeated biotic interchanges between the major landmasses. The early evolution of the main lineages took place in South America, during the Late Cretaceous, and extended to the beginning of the Paleocene, with North American lineages originating from South American relatives. Presence of the clade in the Palearctic region results from multiple exchanges with North America by long-distance dispersal or hopping through landmass chains. Our analyses provide strong support for the amphitropical pattern currently observed in the American continent resulting from at least three periods of possible land connections between the two American landmasses. The last incursion from South to North America is reported in the Miocene, much prior to the well documented Panama Isthmus closure.

## CRediT authorship contribution statement

Kelli Ramos: designed research, data analysis, writing - original draft. Aline Martins: data analysis, writing - review & editing. Gabriel Melo: designed research, data analysis, writing - review & editing.

## Acknowledgments

We are indebted to several colleagues and museum curators for providing access to bee specimens used for DNA extraction and assistance on several aspects of this research: Antonio J. C. Aguiar, Airton T. Carvalho, Arturo Roig-Alsina, Bryan N. Danforth, Danuncia Urban, Eduardo A. B. Almeida, Gabriel A. R. de Paula, Graziele Weiss, John L. Neff, Julia C. Almeida, Luciana Patella, Luis Compagnucci, Olivia Evangelista, Paulo Milet-Pinheiro, Santiago Plischuk, Simone C. Cappellari, and Walter Boeger. Our thanks to Adriana Tiba, Clemens Schlindwein, and Julio Pupim for generously allowing us to use their excellent bee photographs. KSR is very grateful to Daniela M. Takiya, Luis R. R. Faria Jr. and Vitor Kanamura for their invaluable help and companionship in field expeditions. Special thanks to Luisa Ruz for all support during KSR visits in Chile and invaluable support to the development of this research. This work was supported by Coordenação de Aperfeiçoamento de Pessoal de Nível Superior - CAPES (Finance Code 001 to KSR); National Council for Scientific and Technological Development - CNPq (grant 158862/2014-7 to KSR, 304053/2012-0 and 309641/2016-0 to GARM); São Paulo Research Foundation - FAPESP (Grant 2010/17046-5 to KSR); Programa de Capacitação em Taxonomia - PROTAX (Grant numbers CNPq 440574/2015-3 and FAPESP 2016/50378-8 to KSR); CNPq posdoc fellowship to ACM.

## Data availability

Supplementary data are provided in Appendix A. Phylogenetic matrix are provided in Appendix B.

## Notes

### Competing Interest Statement

The authors have declared no competing interest.

